# Deletion of Murine Endothelial *Bag3* Alters the Cellular Proteome Among Organs of Origin

**DOI:** 10.1101/2025.09.24.678094

**Authors:** Zoë S. Terwilliger, Feifei Li, Ananya Pentakota, Tonya N. Zeczycki, Thomas D. Green, Makenzie Kolasa, Nancy C. Edwards, Matthew P. Goldman, Kelsey H. Fisher-Wellman, Joseph M. McClung

## Abstract

BAG3 (Bcl-2 associated athanogene 3) is a multifunctional protein with pleiotropic effects in multiple cell types. Despite our knowledge of its role in cardiovascular disease, its specific role in endothelial cells (ECs) is unknown. The purpose of this study was to identify differences in the EC proteome of multiple tissues before and after cell specific deletion of *Bag3*. We hypothesized that BAG3 loss would uniquely alter the baseline proteome landscape of ECs in each tissue. Cdh5(PAC)-CreERT2;*Bag3*^f/f^ mice received tamoxifen (KO; n=18) or vehicle (WT; n=18) and tissues (brain, heart, lung, and peripheral skeletal muscle-skm) were collected for FACS sorting of ECs from equal numbers of male and female mice at >22 weeks of age, followed by LC-MS/MS label-free proteomics. Initial comparisons of the WT proteomes between tissues revealed differential abundance of EC proteins (p<0.05), including: 379 brain v heart; 325 brain v lung; 501 brain v skm; 354 heart v lung; 107 heart v skm; and 442 lung v skm. Additionally, the EC mitochondrial proteome was unique to each tissue of origin, with significantly (p<0.05) higher proportions dedicated to complexes I, III, IV and V, in heart compared to the other tissues. KO demonstrated the largest effect on skm ECs, increasing 30 proteins (Mybpc2 L2FC=4.81; PFKM L2FC=3.02) and decreasing 65 (CDC42 L2FC=-2.44; Prpf8 L2FC=-2.24). Overall, these results demonstrate that the EC proteome of different tissues is unique and that the loss of BAG3 in these cells differentially alters a small proportion of the EC specific proteome.

## INTRODUCTION

Each organ has different circulatory needs and as such, the endothelium provides specialized services to each location. In cardiovascular diseases, the endothelium serves as a critical physiologic regulator of inflammation, nutrient and gas delivery, and is a crucial site for tipping the scales towards tissue recovery or pathogenesis(1–3). Despite this, our knowledge of the individualized characteristics of local endothelial cells (ECs) lining the inner surface of blood vessels in their host tissues is not thorough. Principle to accurate and strategic therapeutic design is the knowledge of both the differences and similarities of ECs in differing tissue compartments. This is especially important in investigations of gene and target specific manipulations of EC biology in clinical modeling, particularly in the case of genetic targets that have known pathogenic roles in multiple cells where ECs may be affected either directly or indirectly.

Mutations in *BAG3* (Bcl-2 associated athanogene 3) now appear to be one of the most common cause of disease(4), and it’s reported role in multiple cardiovascular diseases makes it an attractive target for investigation in vascular pathologies. BAG3 is an evolutionarily conserved 575 amino acid multifunctional scaffolding protein and co-chaperone with pleiotropic effects in heart and skeletal muscle. It regulates diverse cellular functions and plays critical roles in cancer, skeletal and cardiac muscle biology(5). Mutations in *BAG3* are causally linked to myofibrillar myopathy(6) and dilated cardiomyopathy in humans(7, 8), and loss of *Bag3* in mice causes perinatal lethality due to fulminant muscle myopathy(9). A coding variant in *Bag3* contributes to ischemic muscle necrosis after pre-clinical ischemia, and *Bag3* variant gene therapy targeted to ischemic limb muscles rescues both myopathy and vascular density(10). Previous work has shown that ECs induce BAG3 expression *in vitro* in response to chemopreventative compound exposure and stress,(11) and *in vivo* deletion of EC BAG3 increased oxidative stress and caused pathogenic remodeling after Angiotensin (Ang) II treatment.(12) While additional indirect vascular pathology associated with altered BAG3 function in models of vascular diseases is known(13–16), we cannot currently assign specific importance to BAG3 in EC populations.

In this study we used proteomics to characterize the unique protein signatures of ECs after isolation from four distinct mouse tissues: brain, heart, lung, and peripheral skeletal muscle. We paired this analysis with an EC-specific model of *Bag3* deletion to examine the distinctive changes of this knock-out would have on the proteomes of ECs from these tissues. We hypothesized that each source EC would have a distinct proteomic profile and that EC-specific deletion of *Bag3* would alter the proteome of each tissue in discrete ways. Our results reveal several similarities and differences between the proteomes of ECs directly isolated from different organs, including unique mitochondrial protein signatures. Additionally, we reveal the relative abundances of a small number of unique proteins is altered by the genetic deletion of EC *Bag3* from each organ. Defining a host-tissue’s EC proteome at baseline and in cases of genetic susceptibility fills a critical gap necessary for directional drug development to improve cardiovascular disease outcomes.

## METHODS

### Study Approval and Animal Models

The transgenic *Bag3* knockdown mice were generated by crossing vascular endothelial cell specific, tamoxifen inducible Cdh5(PAC)-CreERT2 male mice (originally developed by Dr. Ralf Adams, London Research Institute, London, UK) with female mice with flox sites flanking exons 2-3 of the *Bag3* gene (*Bag3*^f/f^). *Bag3*^f/f^ 04807 (C57BL/6NTac-Bag3<tm1a(EUCOMM)Hmgu>/H) were purchased from the EMMA biorepository of the International Mouse Phenotyping Consortium (IMPC). The resulting Cdh5(PAC)-CreERT2;*Bag3*^f/-^ male mice from this initial breeding scheme were then crossed with female Bag3^f/f^ mice to generate Cdh5(PAC)-CreERT2;*Bag3*^f/f^ mice. Eight-to-ten-week-old Cdh5(PAC)-CreERT2;*Bag3*^f/f^ mice were injected for five consecutive days with 2mg day-1 of either tamoxifen or vehicle (sunflower seed oil and ethanol) and will be referred to as knockout (KO) and wildtype (WT), respectively. Mice were then aged 20-24 weeks before use. Animals were housed in a climate-controlled environment (23±2) under a 12h light (7:30am-7:30pm)/dark (7:30pm-7:30am) cycle with access to chow and water ad libitum. All mice were bred in-house. All animal experiments adhered to the Guide for the Care and Use of Laboratory Animals from the Institute for Laboratory Animal Research, National Research Council, Washington D.C., National Academy Press, 1996, and any updates. All procedures were approved by the Institutional Animal Care and Use Committees of East Carolina University or Wake Forest University School of Medicine.

### Bag3 Recombination DNA Analysis

Successful recombination was validated in FACS endothelial cells. Tamoxifen induced recombination was confirmed through direct PCR analysis of genomic DNA using a DNA Isolation Kit (QuantaBio ExtractaBio DNA Prep, #95091-025). Primers F – 5’ TCTGAAGATGCACAGGGGTG - 3’ and R – 5’ TTTGGAATTTCACGGCGAGT - 3’; were used to generate an amplicon that was 246bp in length, indicating deletion of the flox region of *Bag3*.

### RNA Isolation and Real Time Polymerase Chain Reaction (RT-PCR)

RNA isolation and qRT-PCR protocols were performed as previously described(17). Total RNA was extracted using Trizol-phenol/chloroform isolation procedures and was reverse-transcribed using Superscript III Reverse Transcriptase and random primers (Invitrogen Inc.). Real-time PCR was performed using a 7500 Real-Time PCR System (Applied Biosystems, Foster City, CA). Relative quantification of *Bag3* mRNA levels was determined using the comparative threshold cycle (ΔΔCT) method using FAM TaqMan® Gene Expression Assays (Applied Biosystems) specific to the given gene run in complex (multiplex) with a VIC-labeled GAPDH control primer.

### Tissue Collection and Cell Preparation

A total of 36 mice (equal numbers male and female) were used for the studies described (18 WT, 18 KO). For each tissue harvest heart, lung, brain, and peripheral limb skeletal muscle tissue were dissected from 6 mice (from each WT and KO) and combined (representing 1 WT or 1 KO, performed 3 separate times for processing and data analysis). The dissected muscle was immediately placed in tubes containing a mixture of Dulbecco’s Modified Eagle’s Medium (DMEM) and PSA. It was then weighed and kept on ice for cell isolation. The dissected muscle was placed on cold petri dishes and manually minced using hemostats and razor blades. From there, three wash cycles, consisting of adding 10 mL of DMEM+PSA, centrifuging at 500xg for 2 minutes, and aspirating the supernatant were done. The minced muscle was transferred to a falcon tube containing 20 mL of warm digest media (20mL DMEM+GlutMax, 2mg/mL Collagenase B, 2.6mg/mL Dispase II, 30% BSA in Saline) and was digested for 30 minutes while rotating. Once the digest was complete, the tube was centrifuged at 500xg for 3 minutes, aspirated, and 5 mL of fetal bovine serum (FBS) was added to stop digestion along with 5 mL of DMEM+PSA.

The resulting pellet was homogenized 5 times using a sterile metal syringe and pipette. After homogenization, the pellet was passed through a 100 µm strainer and centrifuged (400xg, 7 min). After aspirating the supernatant, 1 mL of ACK lysing buffer was added to lyse red blood cells. Once it was added, samples were vortexed for 3 seconds and 10 mL of phosphate buffered saline (PBS) with 2% FBS was immediately added to stop the lysing process. From there, it was centrifuged at (400xg, 7 min) and aspirated before moving on to debris removal. The cell pellet was resuspended in 6.2 mL of 1X PBS and transferred to a 15 ml falcon tube. 1.8 mL of debris removal solution is added and then mixed into the suspension 10 times before adding 4 mL of 1X PBS to create an overlay. Once the overlay was added, it was centrifuged at 3000xg for 10 minutes. After centrifugation was complete, everything except for the cell pellet is aspirated. Then, 13 mL of 1X PBS was added, and the tube was manually inverted three times. It was again centrifuged at 1000xg for 10 minutes and the supernatant was aspirated.

From here, the pellet was resuspended in PBS+2% FBS and centrifuged at 400xg for 7 minutes followed by aspirating the supernatant. The pellet was then resuspended in PBS+0.2% FBS. It was then filtered directly into a fluorescence-activated cell sorting (FACS) tube using a 35µm filter cap.

### Fluorescence-Activated Cell Sorting

Unstained samples were prepared by adding 100µL of the cell stock into a FACS tube with 200µL of PBS+0.2% FBS. The full stain samples were prepared by centrifugation of the remaining stock at 400xg for 7 minutes. The supernatant was then aspirated leaving approximately 90µL in the tube. After the addition of 5µL of FcBlock, the tube was incubated on ice for 15 minutes. The respective antibodies were then added and incubated in the dark for 20 minutes. Then, 3mL of 1x PBS+0.2% FBS was added, and the sample was centrifuged at 500xg for 7 minutes. The tube was decanted and brought to a final volume of 300µL with 1X PBS+0.2% FBS. Antibody capture beads were used as compensation controls. Two drops of each antibody capture bead (positive and negative) were added to a FACS tube. These beads were then equally distributed amongst three tubes to match the three fluorochrome types that were being used (APC, FITC, PE). Next, each antibody that was used with their corresponding amount was added to its respective FACS tube. They were incubated at room temperature in the dark for 15 minutes. 3mL of 1X PBS+2% FBS was added and then the sample was centrifuged at 500xg for 7 minutes. The sample was then aspirated manually, leaving 300µL in the tube. Labeled samples were incubated with CD31 (eBiosciences, #17-0311-82), CD34 (eBiosciences, 11-0341082), and SCA1 (eBiosciences, #12-5981-82). Additionally, 4′,6-diamidino-2-phenylindole (DAPI; D1306, Thermo Fisher) was added to the sample to discern a live cell population. Fluorochrome controls using antibody capture beads are used as compensation controls. Endothelial cells were identified in the heterogenous cell mixture using a set around live cells positive for CD34, CD31, and SCA1. A BD FACSAria^TM^Fusion cell sorter was used for all FACS experiments.

### Sample Preparation and Peptide Isolation for Proteomics

FACS purified ECs were resuspended in buffer (20 mM Tris, pH 7.4, 150 mM NaCl, 1X HALT Protease inhibitor) and lysed via sonication on ice (30% amplitude, 20 sec bursts, 60 sec rest, 4 times). Cold acetone (1:3 v/v) was added to the lysate, the samples were vortexed and placed at −80 C for 2-4 hrs to facilitate protein precipitation. Precipitated proteins were pelleted, washed with neat acetone, and allowed to air dry. Mass spectrometry grade peptides were prepared from the precipitated proteins using the PreOmics iST kit. Briefly, cells were denatured for 10 min in denaturation buffer at 95 °C. Proteins were then alkylated and digested (Tryp/LysC) in one-step for 3 hrs at 37 °C. Digestion was terminated with the addition of stop solution and the entire digestion mixture was added to the provided purification columns. After washing with the provided buffers to remove hydrophobic and hydrophilic contaminates, the bound peptides were eluted with the provided elution buffer in two steps. Peptides were dried by N_2_ stream and resuspended in loading buffer (water: acetonitrile:formic acid, 98%:2%:0.1%). Concentrations were determined by nanodrop (Abs_205_) and adjusting to a final concentration of 0.25 mg/mL.

### LC-MS/MS for Label-Free Proteomics

Peptides were analyzed by nanoLC-MS/MS using an UltiMate 3000 RSLCnano system (ThermoFisher) coupled to a Q Exactive Plus Hybrid Orbitrap mass Spectrometer (ThermoFisher) via nanoeslectrospray ionization. Peptides were separated using an effective linear gradient of 4-35% acetonitrile (0.1% formic acid) over 135 min. For data dependent acquisition in positive mode, MS1 were collected at a resolution of 70,000 with an AGC target of 2×10^5^ ions and a maximum injection time of 100 ms. MS2 spectra were collected on the top 20 most abundant precursor ions with a charge >1 using an isolation window of 1.5 m/z and fixed first mass of 140 m/z. The normalized collision energy for MS2 scans was 30 and were acquired at 17,500 resolution with a maximum injection time of 60 ms, an AGC target of 1×10^5^ and a dynamic exclusion of 30 sec. Proteome Discoverer 2.2 (PDv2.2) was used for raw analysis of data for organ of origin EC comparisons. Default search parameters were oxidation as a variable modification and carbamidomethyl (57.021 Da on C) as a fixed modification, with data searched against both the Uniprot *Mus musculus* reference proteome and the mouse MitoCarta 3.0 database(18). PSMs were filtered to a 1% FDR and grouping to unique peptides was also maintained at a 1% FDR at the peptide level. Strict parsimony was used to group peptides to proteins and filtered to 1% FDR. MS1 precursor intensity was used for peptide quantification. High-confidence master proteins were used to determine mitochondrial enrichment factor (MEF) by quantifying the ratio of mitochondrial protein abundance (identified using the MitoCarta 3.0 database) to total protein abundance, as previously performed(19). Quantification of OxPhos complex subunits expression was determined within each sample by expressing each complex as a percentage of total mitochondrial abundance. FragPipe (v 19.1,(**20, 21**)) was used to for raw analysis of data for KO v WT comparisons with default search parameters for open and Label-Free Quantification Matching Between Runs (LFQ-MBR) workflows. Each sample type (*i.e.* brain, muscle, tissue) were searched individually for each group. An initial open search against the canonical + isoforms Uniprot *Mus musculus* reference proteome (UP000000589, accessed 11/2023 and 1/2024) was used to identify potential post-translational modifications for inclusion in the LFQ-MBR workflow. Precursor m/z tolerance was set to −150 to 500 Da and fragment tolerance was ±20 ppm with 3 missed cleavages for Tryp and Lys-C allowed. Peptide spectrum matches (PSMs) were validated using PeptideProphet and results were filtered at the ion, peptide, and protein level with a 1% false discovery rate (FDR). Based on these initial searches, the following variable modifications were included in the LFQ-MBR analysis: oxidation (+15.5995 Da on Met), deamidation (+0.98401 Da on Gln and Asn), and carbamodiomethyl (+57.025 Da on Cys). For LFQ-MBR analysis, data were search against the canonical Uniprot *Mus musculus* reference proteome (UP000000589, accessed 11/2023 and 1/2024). Precursor ion m/z tolerance was ±20 ppm with 3 missed cleavages for Trypsin/LysC allowed. The search results were filtered by a 1% FDR at the ion, peptide, and protein-level. PSMs were validated using Percolator and label free quantification was carried out using IonQuant(22) Match between runs FDR rate at the ion level was set to 10% for the top 300 runs. Proteins with >95% probability of ID, >2 unique peptides, and in more than 80% of a sample group (*i.e.* 2/3 injections) were considered high confidence IDs and retained for analysis. Intensities were log_2_ transformed, normalized to median intensities of the sample group. Relative abundances for low sampling proteins were determined via normal distribution in Perseus(23).

### Statistics

“The mass spectrometry proteomics data have been deposited to the ProteomeXchange Consortium (http://proteomecentral.proteomexchange.org) #PXD067479 and accession number JPST004017 for jPOST Repository(24, 25). Data for each tissue are provided, including total proteins identified and filtering for ID probability, minimum of 2 peptides and high confidence proteins present in 2/3 injections per group. Spearman correlations were used to identify the relationships between normalized protein abundances and fold changes within organ cell samples between treatments. Significance was set at adjp<0.1, with >1.15fold-change threshold. Quantitative real-time PCR (qRT-PCR), mitochondrial complex proteome comparisons, and mitochondrial enrichment factor (MEF) data was analyzed by PRISM (Version 10.5.0 (673) with p<0.05 considered significant.

## RESULTS

*Endothelial specific deletion of Bag3-* Cdh5(PAC)-CreERT2;*Bag3*f/f mice were bred to generate homozygous deletion littermates (**Figure 1A**). All mice received I.P. injections of either tamoxifen or vehicle at eight weeks of age. WT mice include background matched Cdh5(PAC)- Cre;*Bag3*f/f mice that received vehicle (n=18; **Figure 1B**). KO mice (n=18, 9f/9m) include background matched Cdh5(PAC)-Cre;*Bag3*f/f mice that received tamoxifen. To verify deletion of *Bag3*, genomic recombination was performed on ECs obtained via FACS. Initial gating was performed to remove cell debris from total cell population (**SFigure 1A**). Follow up gating was performed to generate singlet populations, using gating set for only one cell entities (**SFigure 1B**). Next, we performed gating for a live/dead check and CD34^+^ signal (**SFigure 1C**; DAPI^-^ / CD34^+^). Remaining viable CD34^+^ cells are then gated for CD31 and Sca1 (**SFigure 1D**) positivity. Measurements of genomic DNA from FACS-endothelial cells demonstrated that exons 2-3 of the Bag3 gene were deleted (∼246bp) in the endothelial cells of KO mice with no deletion/recombination in the WT mice (∼468bp) (**Figure 1C**). This recombination was specific to the vascular endothelial cells as evidenced by the incomplete recombination observed in the heterogenous mixture of unsorted cells. Genomic recombination was further confirmed by evaluating mRNA expression via qRT-PCR. Reductions in *Bag3* message of the KO group averaged 97.0% (p=0.0135) in comparison to the WT group (**Figure 1D**). Together these data corroborate the specificity and efficiency of *Bag3* in the endothelium in this model.

**Figure 1.**
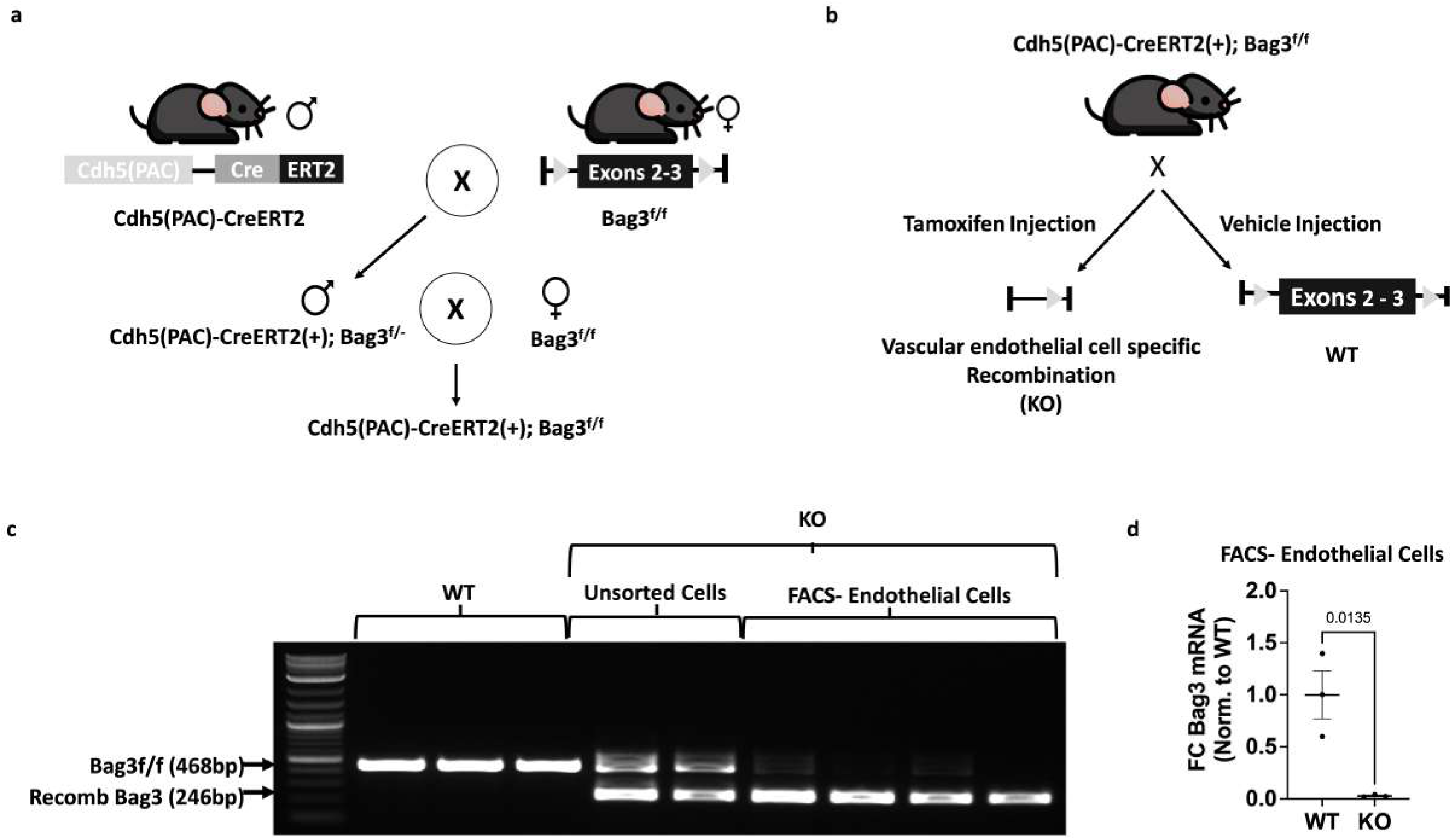
Generation of endothelial cell *Bag3* KO mice. **A.** Cdh5(PAC)-CreERT2;*Bag3*f/f mice were bred to generate homozygous deletion littermates. **B.** All mice received I.P. injections of either tamoxifen (KO; n=18) or vehicle (WT; n=18) at eight weeks of age. C. Fluorescence-activated cell sorting (FACS) was performed to isolate cells from the limb skeletal muscles of WT and KO mice, including unsorted cell populations and an endothelial cell specific fraction. **C**. Recombination polymerase chain reactions were run on genomic DNA from FACS-endothelial cells verifying exons 2-3 of the Bag3 gene were deleted (∼246bp) in the endothelial cells of KO mice with no deletion/recombination in the WT mice (∼468bp). **D**. mRNA expression was verified via qPCR. Values are presented as corrected for 18s, normalized to WT values (Fold Change – FC), means ± SE.

### Organ specific differences in endothelial cell proteomes

To understand how organ specific EC proteomes may differ, we initially investigated the tissue specific proteomes of ECs in WT (Veh) mice. A total of 1484 proteins were identified across tissue ECs (**Figure 2A**). Panther ontology(26) for proteomes between tissues are provided in **Supplemental Figure 1**. Between brain and heart ECs 379 proteins were differentially (p<0.05) expressed, including: 331 increased (ontology classifications predominantly transporter activity and catalytic activity) and 48 decreased (binding and catalytic activity) (**Figure 2B**). Between brain and lung ECs 325 proteins were differentially (p<0.05) expressed, including: 136 increased (transporter activity and catalytic activity) and 189 decreased (binding and catalytic activity) (**Figure 2C**). Between brain and skeletal muscle (SkM) ECs 501 proteins were differentially (p<0.05) expressed, including: 454 increased (translation regulator activity and catalytic activity) and 47 decreased (binding and transporter activity) (**Figure 2D**). Between heart and lung ECs 354 proteins were differentially (p<0.05) expressed, including: 454 increased (binding and catalytic activity) and 47 decreased (binding and transporter activity) (**Figure 2D**). Between heart and skealtal muscle (SkM) ECs 107 proteins were differentially (p<0.05) expressed, including: 94 increased (binding and catalytic activity) and 13 decreased (binding and catalytic activity) (**Figure 2E**). Between lung and SkM ECs 442 proteins were differentially (p<0.05) expressed, including: 416 increased (transporter activity and catalytic activity) and 26 decreased (binding, catalytic activity, and structural molecule activity) (**Figure 2F**).

**Figure 2.**
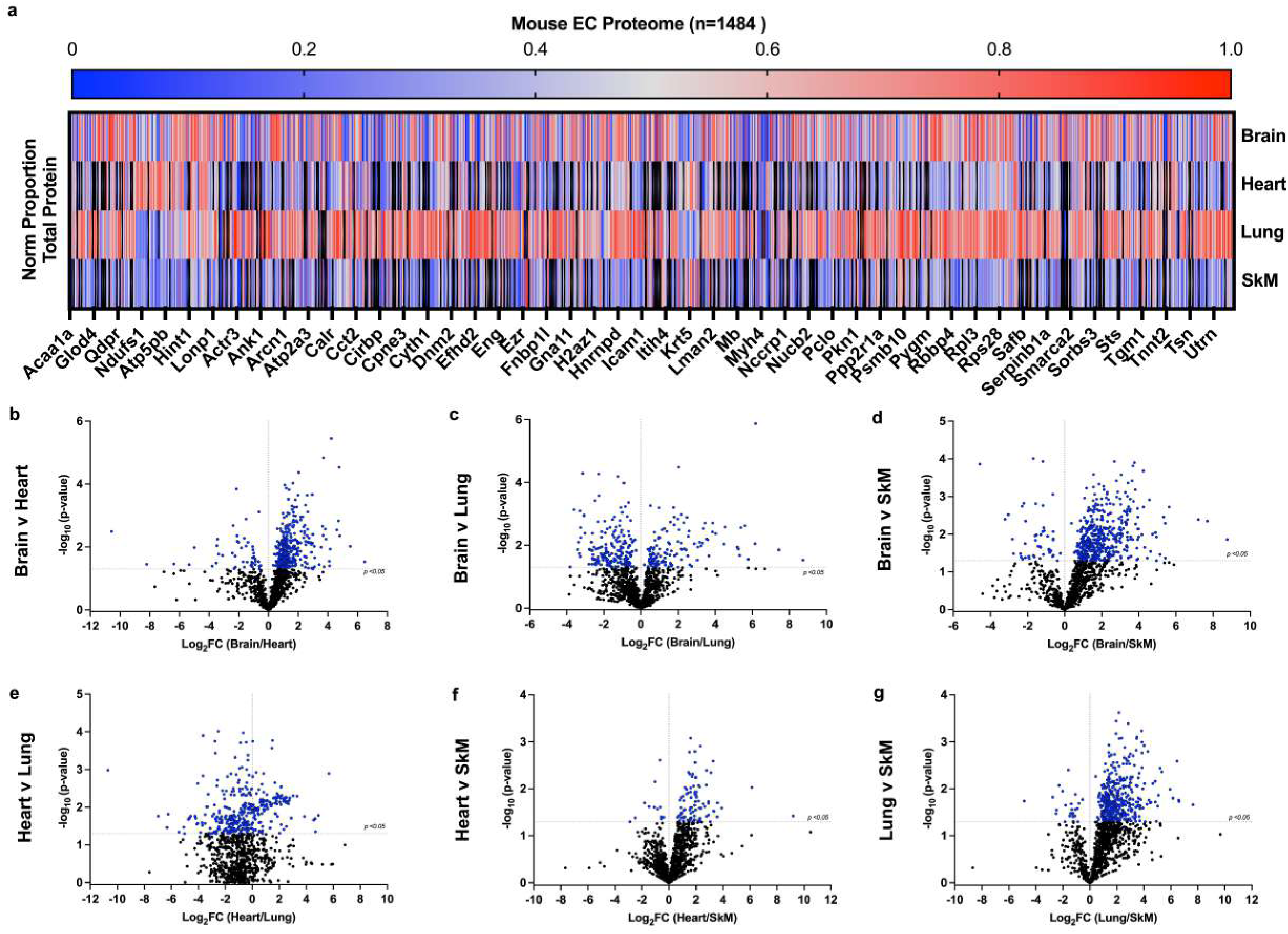
Tissue EC proteomes. **A.** Heat map of EC proteomes from Brain, Heart, Lung, and Skeletal Muscle (SkM), representative of normalized proportion of each total protein identified within tissue. Labels presented for every 30^th^ target (starting at 1). Volcano plots of tissue EC proteome comparisons (y-axis: −log_10_(p-value), x-axis: Log_2_FC). Significance set at adj p<0.05. Brain v Heart (**B**), Brain v Lung (**C**), Brain v SkM (**D**), Heart v Lung (**E**), Heart v SkM (**F**), and Lung v SkM (**G**).

We next examined the EC mitochondrial proteomic profile of each tissue by calculating the summed abundance of all mitochondrial proteins (i.e., MitoCarta 3.0 positive proteins) to total protein abundance. The ‘mitochondrial enrichment factor’ (MEF; **Figure 3A**) was uniquely reduced in the SkM ECs versus either Brain or Heart ECs and reflects cells with much smaller proportional mitochondrial proteins in their ECs. Additionally, the EC mitochondrial proteome was unique in each tissue of origin, with significantly (p<0.05) higher proportions of the mitochondrial proteome dedicated to complexes I, III, IV and V in heart ECs compared to the other tissues (**Figure 3B**). The overall percentage of the MitoCarta proteome apportioned to complexes I, II, III, and IV in Brain, Heart, Lung, and SkM ECs, and Complex V in Lung and SkM are much lower than the values reported for other cell types(27). Heat mapping of the MitoCarta proteome across tissues reveals strong proportional mitochondrial proteins in the Brain and Heart ECs, including predominant expression of some targets (Gpd2, Eci2, Pccb, Phb1) in Brain ECs (**Figure 3B**). Overall, this collective data provides important baseline insight into the EC proteome across multiple tissues of origin. What follows are the individual tissue EC proteome comparisons after deletion of Bag3.

**Figure 3.**
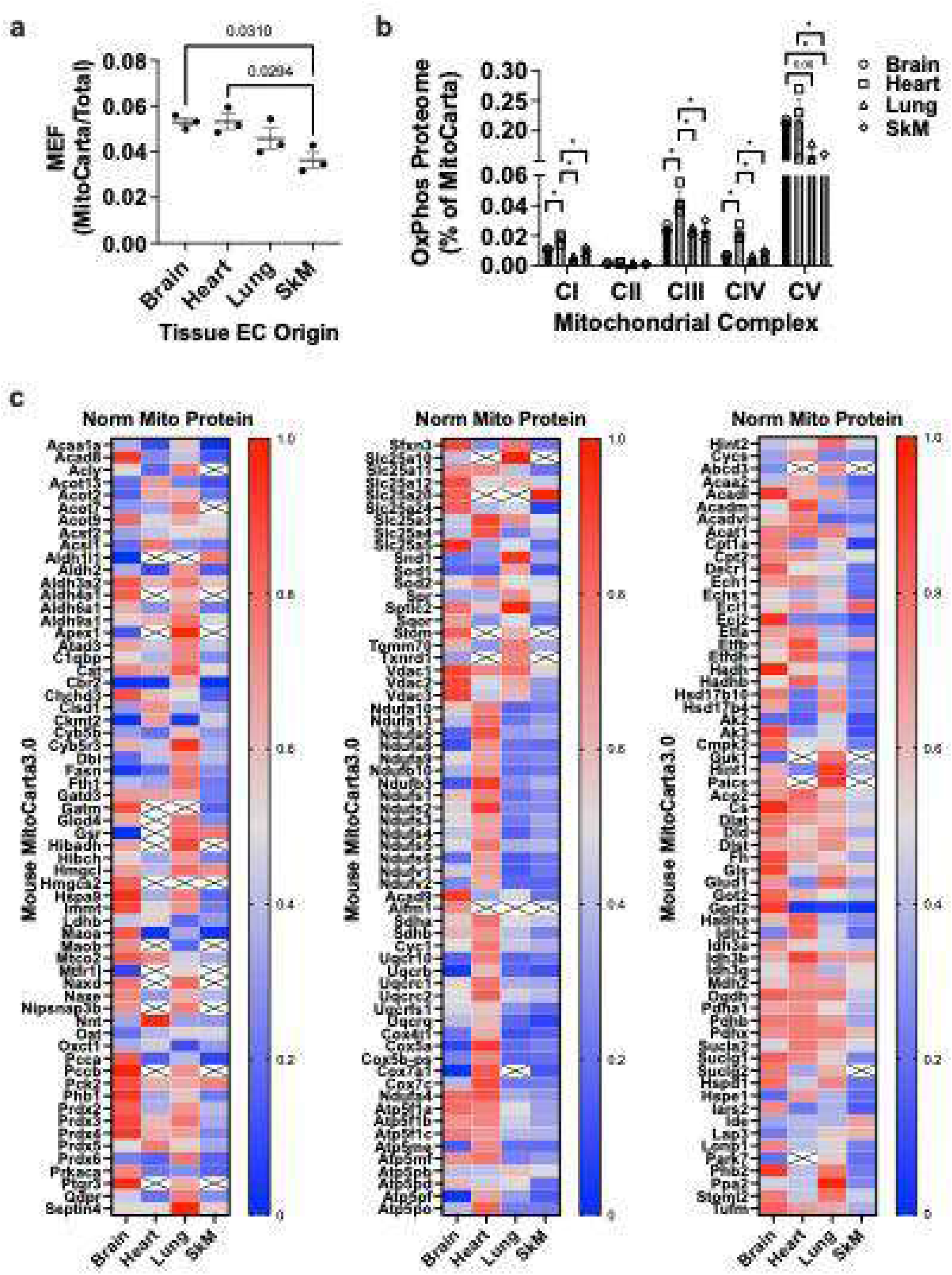
Tissue EC mitochondrial proteome. EC proteomic data from Brain, Heart, Lung, and skeletal muscle (SkM) was sorted for mitochondrial proteins (MitoCarta). **A.** Mitochondrial enrichment factors (MEF) per tissue ECs, derived from the ratio of mitochondrial to total protein abundances. **B.** Identified mitochondrial proteome components were sorted by mitochondrial complex (I, II, III, IV, and V) and presented as the % of mitochondrial proteome within each tissue. Values are presented as means ± SE, * p<0.05 or labeled (0.08). **C.** Heat map of EC mitochondrial proteomes from Brain, Heart, Lung, and SkM. Labels presented for each target, representative of normalized proportion of each total protein identified within tissue (n=3/tissue). X indicates blank values.

### Bag3 deletion in the Brain endothelium

On average 400,000 ECs were isolated per sample (*n*=3, 6 mice pooled/sample) after FACS in vehicle treated (WT) Brain tissues and 600,000 ECs were isolated per sample (*n*=3) in tamoxifen treated (KO) brain tissues. A total of 1893 proteins were identified in the FACS brain tissue EC population. Filtering revealed 1798 proteins with ID probability >0.955. There were 1397 proteins with a minimum of 2 peptides identified and 1377 proteins with high confidence (in at least 2/3 injections per sample). Spearman correlation between WT and KO Log2 Abundances revealed a strong relationship between WT and KO protein abundances (R^2^ =0.9308 and p<0.0001; **Figure 4A**). Volcano plots of all proteins (164 total) in KO ECs (p<0.1) v WT (**Figure 4B**) demonstrates a pattern of increased protein abundances in KO ECs. Corresponding graphical identification of 72 increased proteins and 43 decreased (**Figure 4C,D**) proteins (*p*<0.1, ≥ 1.15FC) in KO ECs is provided.

**Figure 4.**
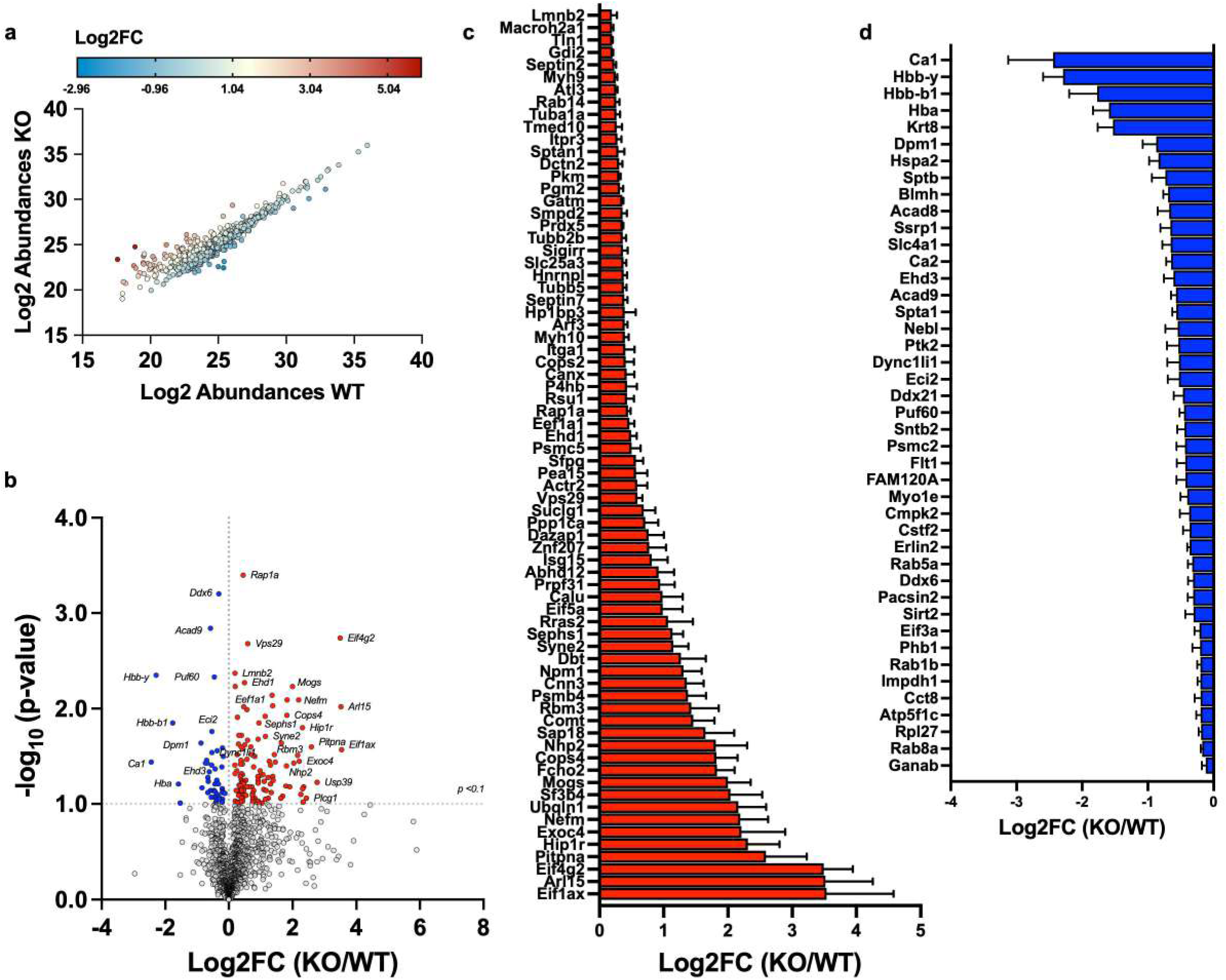
*Bag3*KO effects on the Brain EC proteome. ECs were isolated from WT (n=3) and KO (n=3) Brain samples for proteomics. A total of 1377 proteins were identified with high confidence across the samples. **A**. Spearman correlation between WT and KO Log_2_ Protein Abundances revealed a strong relationship between WT and KO proteins (R^2^=0.9308 and p<0.0001). **B**. Volcano plot of all proteins, colors designate proteins with significant abundance decreases (blue) and increases (red) in KO ECs (p<0.1, 164 total proteins) compared to WT. **C.** Corresponding graphical identification of 72 increased proteins and (**D**) 43 decreased proteins (*p*<0.1, ≥ 1.15FC) in KO ECs. Values are presented as means ± SEM.

### Bag3 deletion in the Heart endothelium

Across the vehicle treated (WT) heart tissues on average 630,000 ECs were isolated per sample (*n*=3) after FACS. Across the tamoxifen treated brain tissues on average 540,000 ECs were isolated per sample (*n*=3). A total of 944 proteins were identified in the FACS heart tissue EC population. Filtering revealed 833 proteins with ID probability >0.955. There were 674 proteins with a minimum of 2 peptides identified and 673 proteins with high confidence (in at least 2/3 injections per sample) inference identified in total. Spearman correlation between WT and KO Log2 Abundances revealed a strong relationship between groups (r=.9653 and p<0.0001; **Figure 5A**). Volcano plotting demonstrates 46 proteins *p*<0.1 between WT and KO ECs (**Figure 5B**). Corresponding graphical identification of 33 increased proteins (*p*<0.1, ≥ 1.15FC) and 7 decreased proteins in KO ECs (**Figure 5C**).

**Figure 5.**
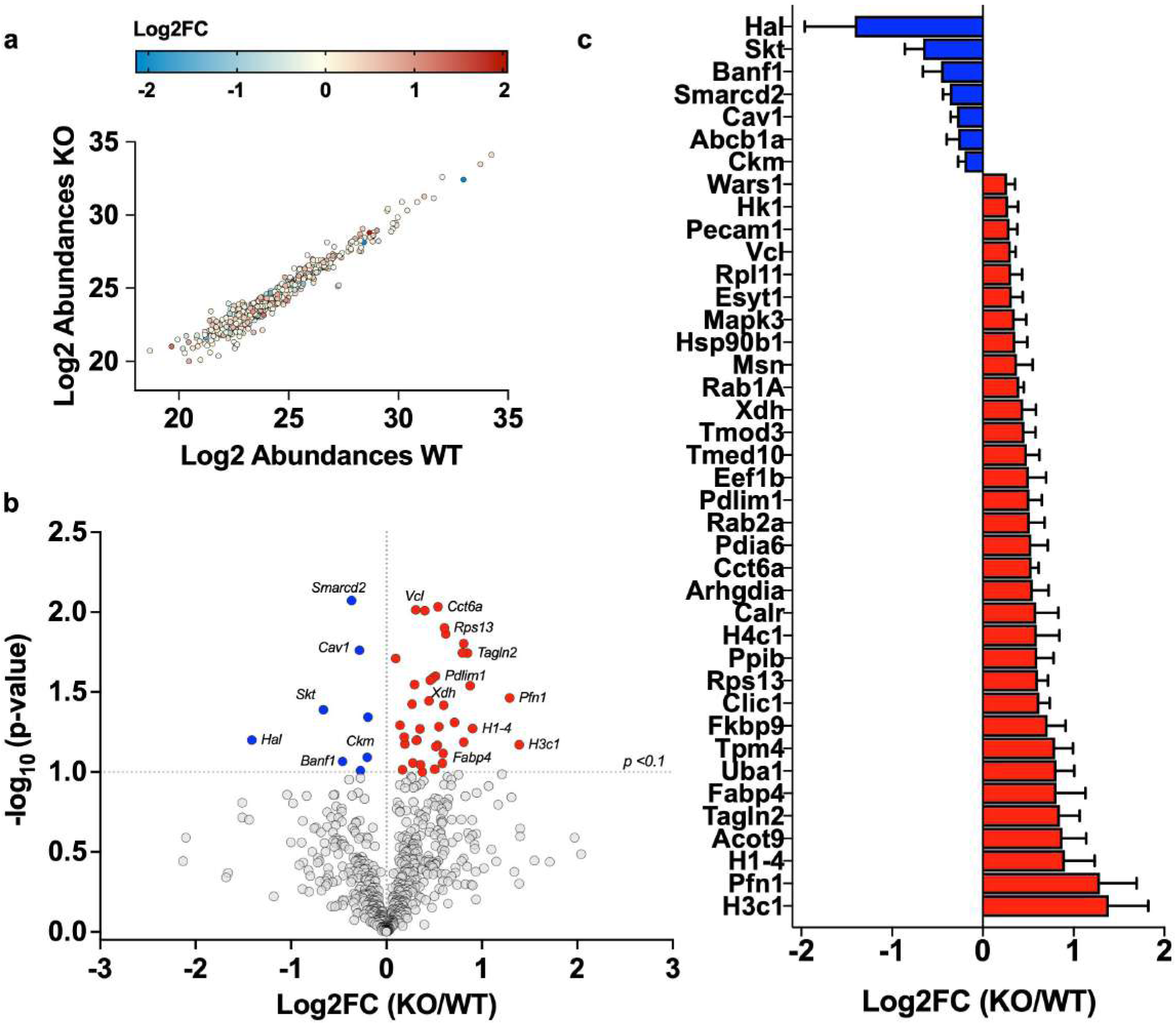
*Bag3*KO effects on the Heart EC proteome. ECs were isolated from WT (n=3) and KO (n=3) Heart samples for proteomics. A total of 673 proteins were identified with high confidence across the samples. **A**. Spearman correlation between WT and KO Log2 Abundances revealed a strong relationship between WT and KO proteins (r=.9653 and p<0.0001). **B**. Volcano plotting of all proteins, colors designate decreased (blue) and increased (red) proteins (46 total) in KO ECs (p<0.1) v WT. **C.** Corresponding graphical identification of 33 increased proteins and 7 decreased proteins (*p*<0.1, ≥ 1.15FC) in KO ECs.

### Bag3 deletion in the Lung endothelium

Across the vehicle treated (WT) lung tissues on average 366,000 ECs were isolated per sample (*n*=3) after FACS. Across the tamoxifen treated brain tissues on average 550,000 ECs were isolated per sample (*n*=3). A total of 1816 proteins were identified in the FACS lung tissue EC population. Filtering revealed 1676 proteins with ID probability >0.955. There were 1368 proteins with a minimum of 2 peptides identified and 1352 proteins with high confidence (in at least 2/3 injections per sample) inference identified in total. Spearman correlation between WT and KO Log2 Abundances revealed a strong relationship between groups (r=.9618 and p<0.0001; **Figure 6A**). Volcano plotting revealed 77 proteins *p*<0.1 between WT and KO ECs (**Figure 6B**). Graphical identification of 34 increased proteins (*p*<0.1, ≥ 1.15FC) and 30 decreased proteins in KO ECs is provided in **Figure 6C**.

**Figure 6.**
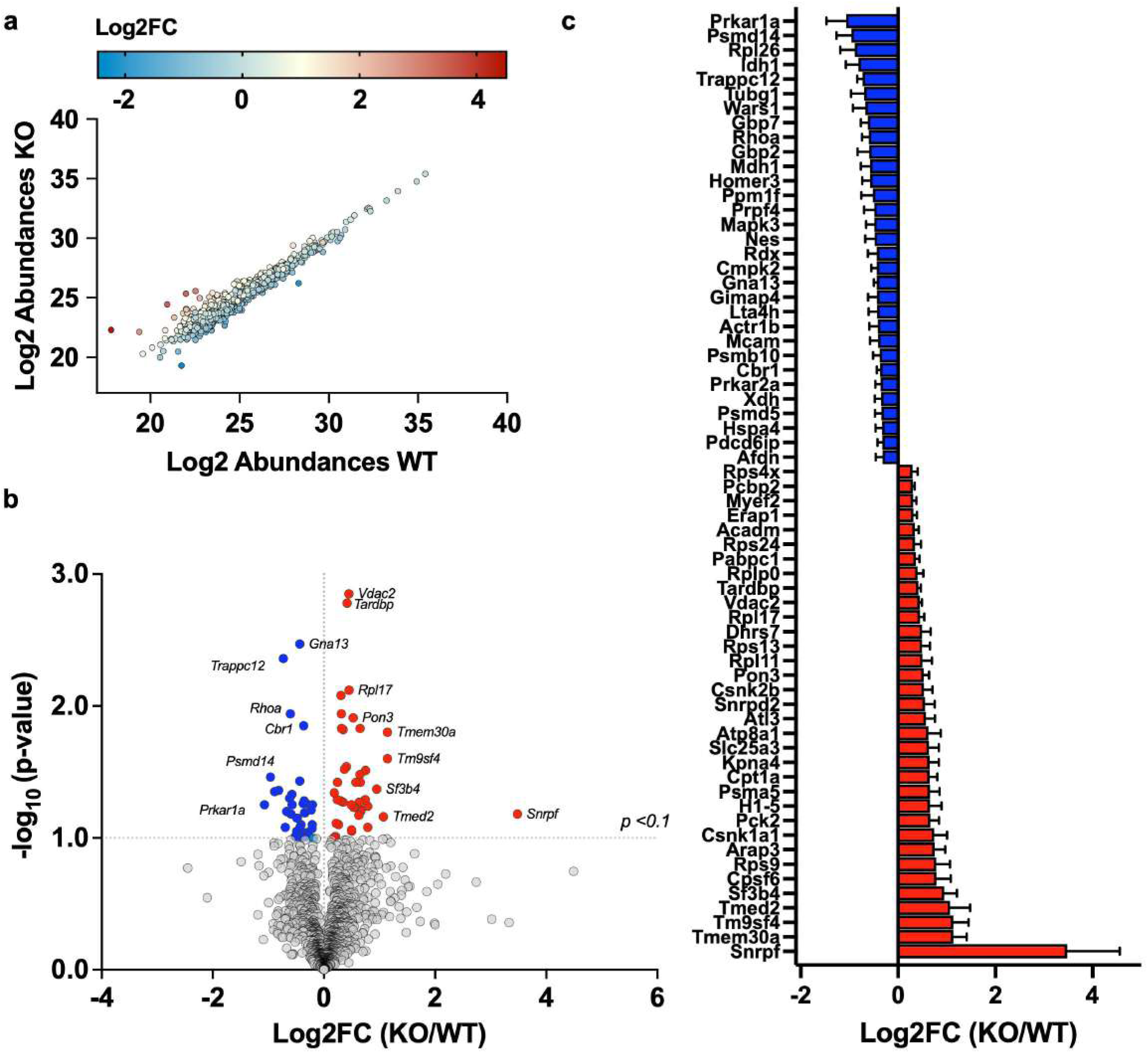
*Bag3*KO effects on the Lung EC proteome. ECs were isolated from WT (n=3) and KO (n=3) Lung samples for proteomics. A total of 1352 proteins were identified with high confidence across the samples. **A**. Spearman correlation between WT and KO Log2 Abundances revealed a strong relationship between WT and KO proteins (r=.9618 and p<0.0001). **B**. Volcano plotting of all proteins, colors designate decreased (blue) and increased (red) proteins (77 total) in KO ECs (p<0.1) v WT. **C.** Corresponding graphical identification of 34 increased proteins and 30 decreased proteins (*p*<0.1, ≥ 1.15FC) in KO ECs.

### Bag3 deletion in the peripheral skeletal muscle endothelium

Across the vehicle treated (WT) skeletal muscle tissues on average 500,000 ECs were isolated per sample (*n*=3) after FACS. Across the tamoxifen treated brain tissues on average 455,000 ECs were isolated per sample (*n*=3). A total of 687 proteins were identified in the FACS skeletal muscle tissue EC population. Filtering revealed 600 proteins with ID probability >0.955. There were 493 proteins with a minimum of 2 peptides identified and 481 proteins with high confidence (in at least 2/3 injections per sample) inference identified in total. Spearman correlation between WT and KO Log2 Abundances revealed a strong relationship between groups (r=.8317 and p<0.0001) (**Figure 7A**). Volcano plotting revealed 96 proteins *p*<0.1 between WT and KO ECs (**Figure 7B**), including 31 proteins >1.15-fold increased (*p*<0.01) and an additional 65>1.15-fold decreased (*p*<0.01). Graphical identification of 31 increased proteins and 65 decreased proteins (*p*<0.1, ≥ 1.15FC; **Figure 7C,D**) in KO ECs.

**Figure 7.**
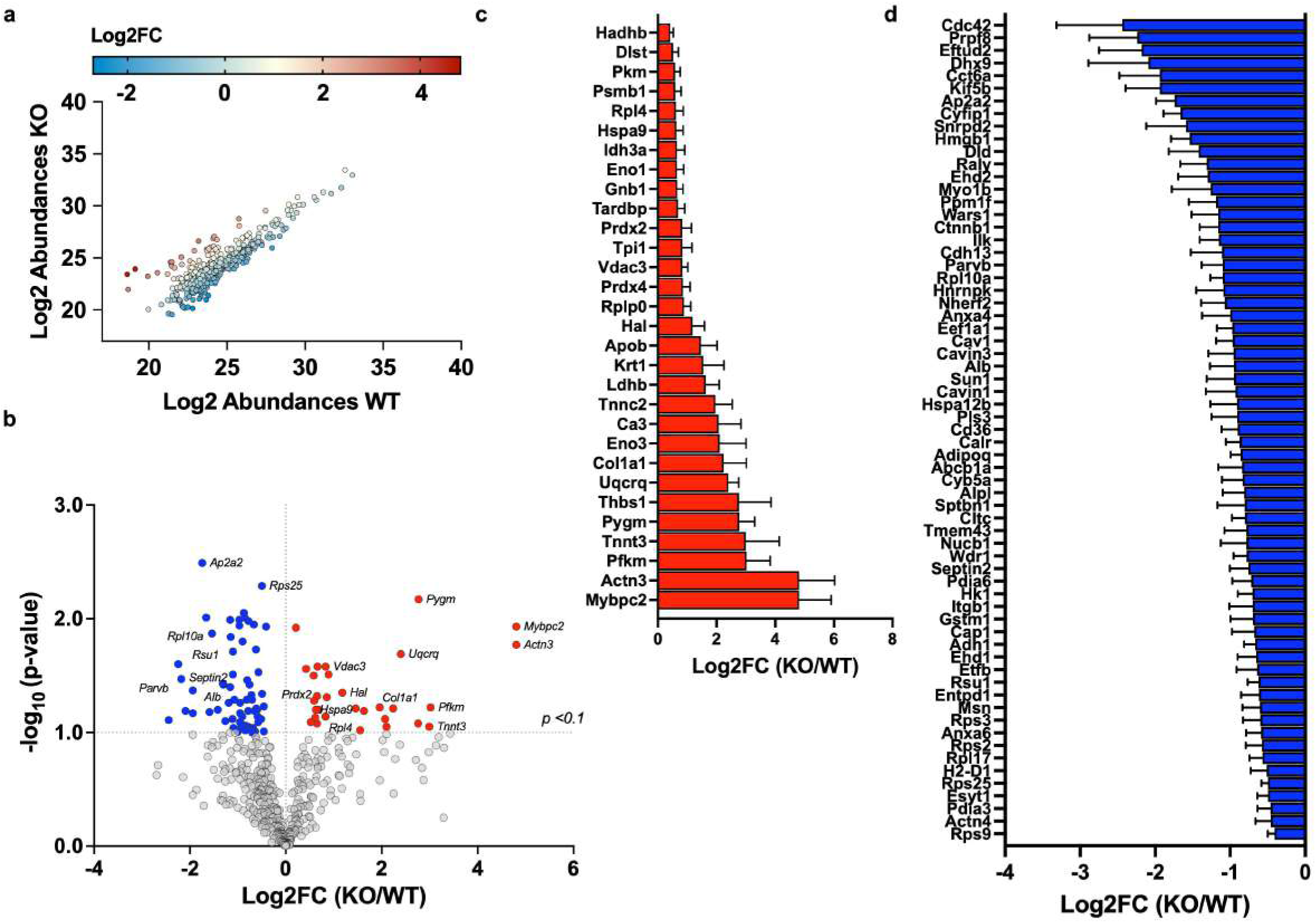
*Bag3*KO effects on the SkM EC proteome. ECs were isolated from WT (n=3) and KO (n=3) SkM samples for proteomics. A total of 481 proteins were identified with high confidence across the samples. **A**. Spearman correlation between WT and KO Log2 Abundances revealed a strong relationship between WT and KO proteins (r=.9317 and p<0.0001). **B**. Volcano plotting of all proteins, colors designate decreased (blue) and increased (red) proteins (96 total) in KO ECs (p<0.1) v WT. **C.** Corresponding graphical identification of 30 increased proteins and (**D**) 65 decreased proteins (*p*<0.1, ≥ 1.15FC) in KO ECs.

### Common targets of EC Bag3KO across organs

We lastly compiled a complete list of proteins showing significantly different abundances in (*p*<0.1) WT and KO ECs across all 4 organs to look for commonalities. Among the 383 proteins found to be affected by the deletion of BAG3, there were no proteins which showed differential abundances in 3 or more organs of origin. Only 14 identified targets were changed by KO between 2 organs (similarly or differentially) (**Figure 8**). The largest number of target similarities were found between the Brain and SkM ECs after KO (5 total, 2 similar directionally). The least number were between the Lung and Heart (1 total, similar directionally). Bag3KO ECs from Lung and Brain directionally matched 3/3 (increased) targets.

**Figure 8.**
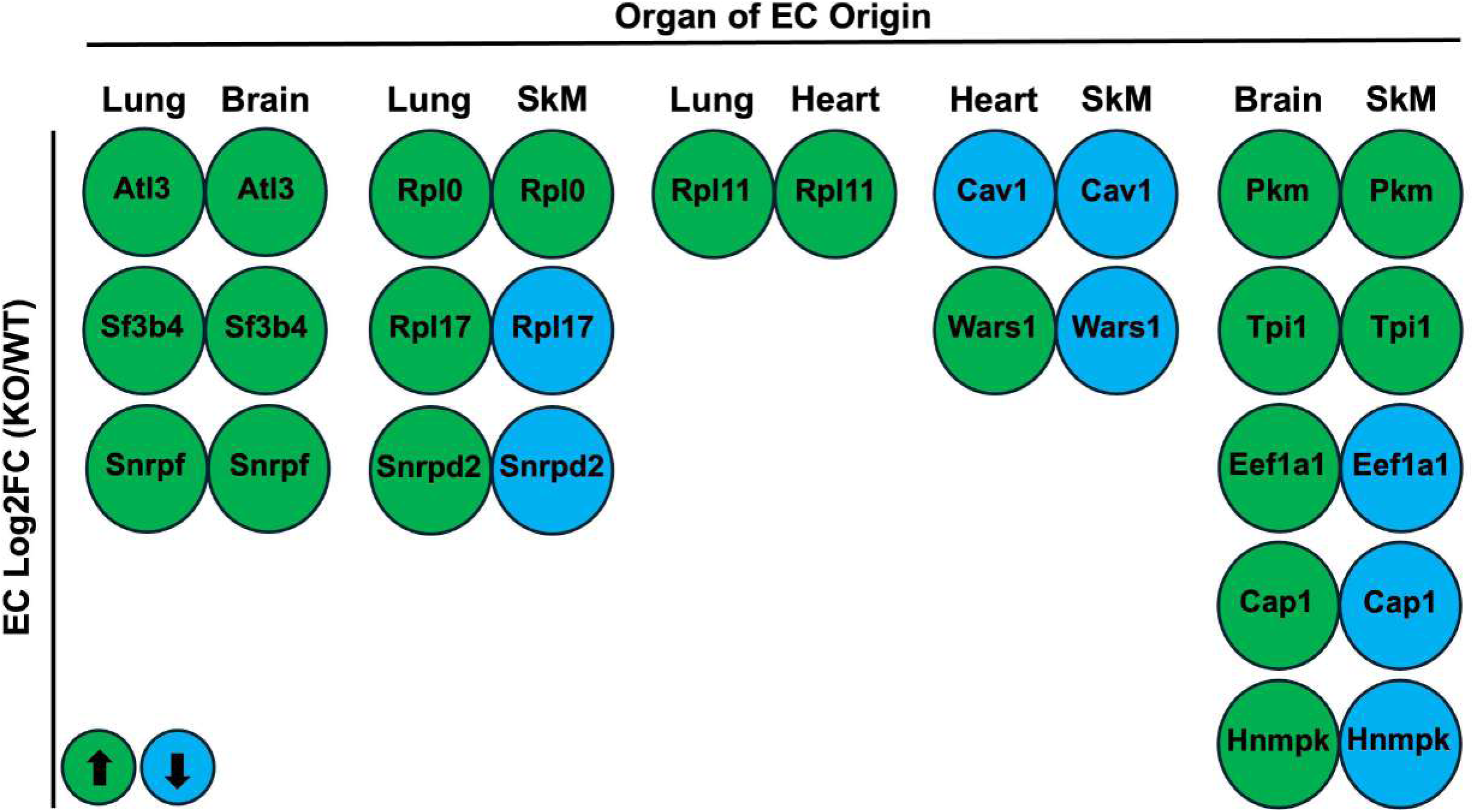
Common targets of EC *Bag3*KO across organs. All proteins significantly (*p*<0.1) altered by *Bag3*KO (KO/WT; Log2FC; n=383) across all 4 organs of EC isolation were analyzed for common targets and directionality (green/increase; blue decreased). Only 14 identified targets were altered by KO between 2 organs.

## DISCUSSION

In this study, we used LC-MS/MS for label-free proteomics on ECs isolated from 4 different organs from mice with or without genetic deletion of Bag3. We analyzed the proteomics data in two formats, first as WT EC between organs and second, within organs between mice with tamoxifen induced KO of Bag3. Our proteomic analysis demonstrates a unique proteomic profile of ECs specific to each tissue type. While we are not the first to describe unique EC proteomes in cells harvested from different tissues(28–30) our approach provides a unique perspective on primary ECs naïve to any *in vitro* expansion. Brain ECs provided the most contrasting cellular proteomes when compared with other tissues, followed by cells isolated from the Lung. ECs isolated from the brain had widely variable proportions of proteins when compared with the other tissues examined. Proteins like Histidine triad nucleotide binding protein 1 (HINT1), a tumor suppressor, and Lon peptidase 1, an essential mitochondrial protease responsible for protein quality control and response to cellular stress(31–34), were enriched in brain ECs compared to other organs. Myosin heavy chain 4 (MYH4), a key protein in muscle contraction, and chaperonin containing TCP1 subunit 2 (CCT2), a chaperone protein that assists in correct formation of nascent proteins, showed decreased abundances in brain ECs. The most similar EC proteomes belonged to Heart and SkM. Interestingly, the tissue EC proteomes included differential mitochondrial protein compositions, with SkM endothelial cells possessing the lowest enrichment of mitochondrial proteins. We also identified unique mitochondrial complex specific proteins between the ECs from each tissue, providing insight into differential organelle organization/priority within each parent organ.

Observed changes in the EC specific mitochondrial proteins in different organs of origin is interesting, particularly in the context of EC bioenergetics. ECs were long believed as simple conduits that rely on glycolytic ATP production for angiogenesis(35–37). Recently, critical roles for EC mitochondria have been identified, including: fatty acid oxidation mediated EC progenitor cell regulation(38); deficiencies in EC substrate oxidation causing a pathologic endothelial mesenchymal transition(39); and mitochondrial substrate oxidation being required for EC dNTP synthesis, vessel sprouting, and proliferation(40). The reported values for ‘mitochondrial enrichment factor’ (MEF) across all the tissue ECs are smaller than values reported for other whole tissue isolates (27, 41, 42). This is perhaps reflective of a lesser need for mitochondrial protein density in the endothelium versus other cell types. Similar to reductions in MEF, the overall % of the MitoCarta proteome apportioned to complexes I, II, III, and IV in Brain, Heart, Lung, and SkM ECs, and Complex V in Lung and SkM are much lower than the values reported for other cell types (42, 43). Finding that SkM ECs possessed uniquely reduced MEF values and correspondingly smaller proportional proteins affiliated with several mitochondrial complexes is an exciting finding and demonstrates a specific characteristic of these EC populations that could impact EC biology under stress, including vascular diseases of the periphery. Our data demonstrates that under normal conditions the ECs from more metabolically active tissues such as the Brain(44–49) and Heart(43, 48–53) contain more mitochondrial proteins. ECs isolated from the heart had the highest expression of Ndufs1 and Atp5pb, two mitochondrial proteins responsible for creating the NADH:ubiquinone oxidoreductase core subunit S1 and the b subunit of the proton channel within ATP synthase, respectively. Higher metabolic capacity in the vascular ECs in these tissues could be related to the proportion of blood flow that these organs normally demand(54–58), but points to a unique characteristic of these specific EC populations that warrants further investigation.

Deletion of *Bag3* had the largest effect on the Brain EC proteome, with 164 proteins differentially expressed. The most striking decreases in Brain ECs were related to the hemoglobin associated proteins, including HBB-y (gamma globulin-subunit), HBB-b1 (beta globulin-subunit), and Hba (alpha globulin-subunit). All three proteins were significantly downregulated with KO. Hba is known to regulate vascular tone and function in resistance arteries.(59) Gamma and beta globulin subunits have not been deescribed in mouse resistance vessels, although their presence in the sorted EC proteomes here may support the assertion that their expression in other endothelial populations, including exchange vessels such as capillaries, varies.(60) A universal downregulation indicates across multiple vessel endothelial cell types that hemoglobin stabilization protein expression is a unique target of *Bag3*KO. Among the proteins increased in Brain derived ECs, several increases were observed targets associated with protein synthesis, including large increases in EIF1ax (eukaryotic translation initiation factor 1A, X linked), Eif3L (eukaryotic translation initiation factor 3, subunit L), and EIF4g2 (eukaryotic translation initiation factor 4 gamma 2), and smaller increases in proteins associated with translational elongation Eif5a (eukaryotic translation initiation factor 5A) and Eef1a1 (eukaryotic translation elongation factor 1 alpha 1). Increases in both initiation and elongation factors for protein synthesis would indicate that regulation of protein synthesis is uniquely affected in Brain ECs with *Bag3*KO. BAG3 is a known regulator of proteostasis and synthesis in striated muscles.(61, 62) The initiation factor proteins are linked to protein synthesis via translational initiation,(63) although EIF4G2 has also been reported to function as a repressor by driving the formation of inactive complexes(64).

KO also had a large effect on the proteomic signature of skeletal muscle ECs. Mybpc2 (myosin binding protein C2) and Pygm (glycogen myelophosphorylase) saw the largest fold increases at the lowest p-values in the muscle ECs. Increased Mybpc2 is of interest due to its known role in angiogenesis.(65) Increased Pygm, paired with observed increases in (3.0 Log2FC) in Pfkm (phosphofructokinase) and Pkm (0.58 Log2FC) indicate a unique upregulation of proteins related to glycolysis in KO ECs. Pygm plays a role in breaking down glycogen to form glucose, Pfkm is one of the most highly regulated, ATP-requiring steps in glycolysis and catalyzes the phosphorylation of fructose-6-phosphate to fructose-1,6-bisphosphate, and Pkm is specifically involved in the conversion of phosphoenolpyruvate to pyruvate. KO also upregulated Enolase 1 (ENO1) and Eolase 3 (ENO3), which are enzymes key to promoting glycolysis(66, 67). KO downregulated proteins in SkM ECs included Cdc42 (cell division cycle 42), a key protein involved in maintaining the EC barrier and vessel permeability(68) and Ehd2 (Epsin-homology domain protein 2), a protein involved in sprouting regulation during vessel sprouting.(69) Upregulation of proteins related to glycolytic flux and downregulation of proteins associated with vessel barrier integrity and aberrant blood vessel sprouting is an ominous sign for vasculature and could indicate changes related to dysfunctional vessel angiogenesis or leaky vessels.(70, 71) Further studies are needed to provide context to these baseline changes during vascular stressors.

*Bag3* mutations or KO have profound impacts on the Heart,(61, 62, 72–75) although most studies are focused specifically on cardiac muscle. We were initially surprised that the cardiac EC proteome was the least altered cellular proteome after KO in this study, including only 3 proteins with +/- > 1.29 Log2FC differences. The overwhelming majority of differences were increases in EC proteins, including PECAM1 (Platelet and Endothelial Cell Adhesion Molecule 1; CD31) and Calr (calreticulin), both of which play known roles in endothelial cell function.(76, 77) Additionally Pfn1 (profilin-1) protein was increased in KO ECs, which has reported significant impacts on angiogenesis and EC biology(78, 79). Collectively, increasing cell adhesion and cell structural dynamics related proteins such as these supports the possibility of structural changes to the EC barrier in the heart vasculature.

Like the Heart ECs, Lung ECs displayed a somewhat muted cellular proteome change with the BAG3 KO. While the Lung saw more total targets significantly altered, by the same metric utilized above to describe the heart endothelium (>1.29Log2FC) SNRPF (Small Nuclear Ribonucleoprotein Polypeptide F) was the only target with large differences in expression post-KO. SNRPF is a core spliceosome protein and is critical to pre-mRNA processing for protein synthesis. Circulatory SNRPF has been identified in a quantitative trait locus for type 2 diabetes,(80) although any direct molecular connectivity in the context of Bag3KO in or outside of ECs requires additional work. Lung ECs displayed a pattern of downregulation of proteins related to guanine nucleotides, including Gbp2 (Guanylate Binding Protein 2), Gnb4 (guanine nucleotide-binding protein subunit beta-4), Gbp7 (Guanylate Binding Protein 7), and RhoA (Ras homolog family member A). These proteins collectively control many cellular processes and intracellular signaling in ECs (81–83), but are well known to regulate cytoskeletal arrangement and barrier integrity between the endothelium and angiogenesis(84, 85), which indicate a pattern of reduced proteins related to vessel stability.

In looking across the organ proteome changes with Bag3KO, there were no observed universally decreased or increased proteins. No protein was altered in more than two organs after KO, and only 8 proteins were directionally (increased or decreased) similarly in 2 organs simultaneously (Pkm, Rplp0, Sf3b4, Rpl11, Snrpf, Tpi1, Cav1, and Atl3). While we didn’t initially hypothesize that changes would occur in this fashion, it provides further evidence of the heterogeneity of the collective endothelial cell populations. This idea is well represented in the differences between the muscle organs, heart and SkM. Cardiac and SkM are two tissues known to be severely affected by changes in BAG3 expression and function(10, 61, 72, 86–94) and could reasonably be hypothesized to have their ECs affected most similarly with KO. While there were a few noticeable differences in the proteomes of these ECs in WT mice, each organ had uniquely different responses. Cav1 (caveolin-1) was the only protein similarly decreased in ECs isolated from these tissues after KO. Caveolin is a protein involved in creating membrane invaginations in cells that allow for lipid homeostasis, transcriptional regulation, signal transduction, and mechanical protection against the vigorous swelling and stress created by contracting straited muscle.(95)

It has been well reported that ECs from varying vessel types (artery v. capillary v. vein) are different and are required to perform specialized functions. Arteries, for example, have ECs that are surrounded by smooth muscle and are responsible for controlling vascular tone while ECs from post-capillary venules express receptors for histamine and serotonin and lack abundant tight junctions which makes them highly permeable during inflammatory conditions(96, 97). Similarly, each of the four tissues investigated in this study perform vastly different functions which likely affects their EC density, specialization, and quantity(97, 98). Because of this, it can be assumed that the described data reflect the intrinsic heterogeneity of each individual tissue.

Previous studies investigated the differences in EC proteome of different tissues. Groten et al. demonstrated that PLEKHA5 and CD109 were associated with aorta, SAPCD2 and TFPI2 with brain, PHLDA1 and SPOCK1 with cardiac, VAMP8 with lung, and ERBB2 with kidney in human tissues(28). When ECs of liver, lung, and kidney were exposed to inflammatory cytokines an upregulation of proteins involved in DNA metabolism, defense/immunity, gene specific transcriptional regulation, protein modification, and anti-apoptosis occurred(30). Simultaneously, a downregulation of proteins involved in cytoskeletal (non-motor actin binding), extracellular matrix, coagulation, apoptosis, cell cycle, protein binding, angiogenesis, cell proliferation, metabolite interconversion, and calcium binding occurred(30). These studies highlight the adaptability of the EC proteome from different organs to disease states and physiologic stress. The work performed herein sets the stage for future work in vascular diseases that involve EC BAG3.

## Statement of Ethics

This animal study was approved by the institution. All experiments were performed in accordance with the *Guide for the Care of Laboratory Animals* published by the National Institutes of Health (NIH Guide, 8^th^ edition, 2011).

## Conflict of Interest

The authors have conflicts of interest to declare.

### Funding Sources

This work was supported in part by NIH grant R01HL125695 (JMM).

## Author Contributions

ZST, FL, and MK performed all *in vivo* work and proteomics preparation. AP and TDG conducted the FACS. KHFW, TNZ, and TDG provided supervision and interpretation of proteomics results. NCE and MPG assisted in manuscript preparation. JMM provided supervision and interpretation of the results, assisted with preparation of manuscript, and raised funds for the study.

### Author Statement

JMM lab (JMM, ZST, FL, TDG) moved from Brody School of Medicine at East Carolina University to Wake Forest University School of Medicine during the performance of the work described herein.

## Figure Acknowledgements

’Mouse Image: Flaticon.com’. This image has been designed using resources from Flaticon.com

## Data Availability

All data generated or analyzed are included in this article and online supplementary materials. Additional inquiries can be directed to the corresponding author.

---

**Supplemental Figure 1.**
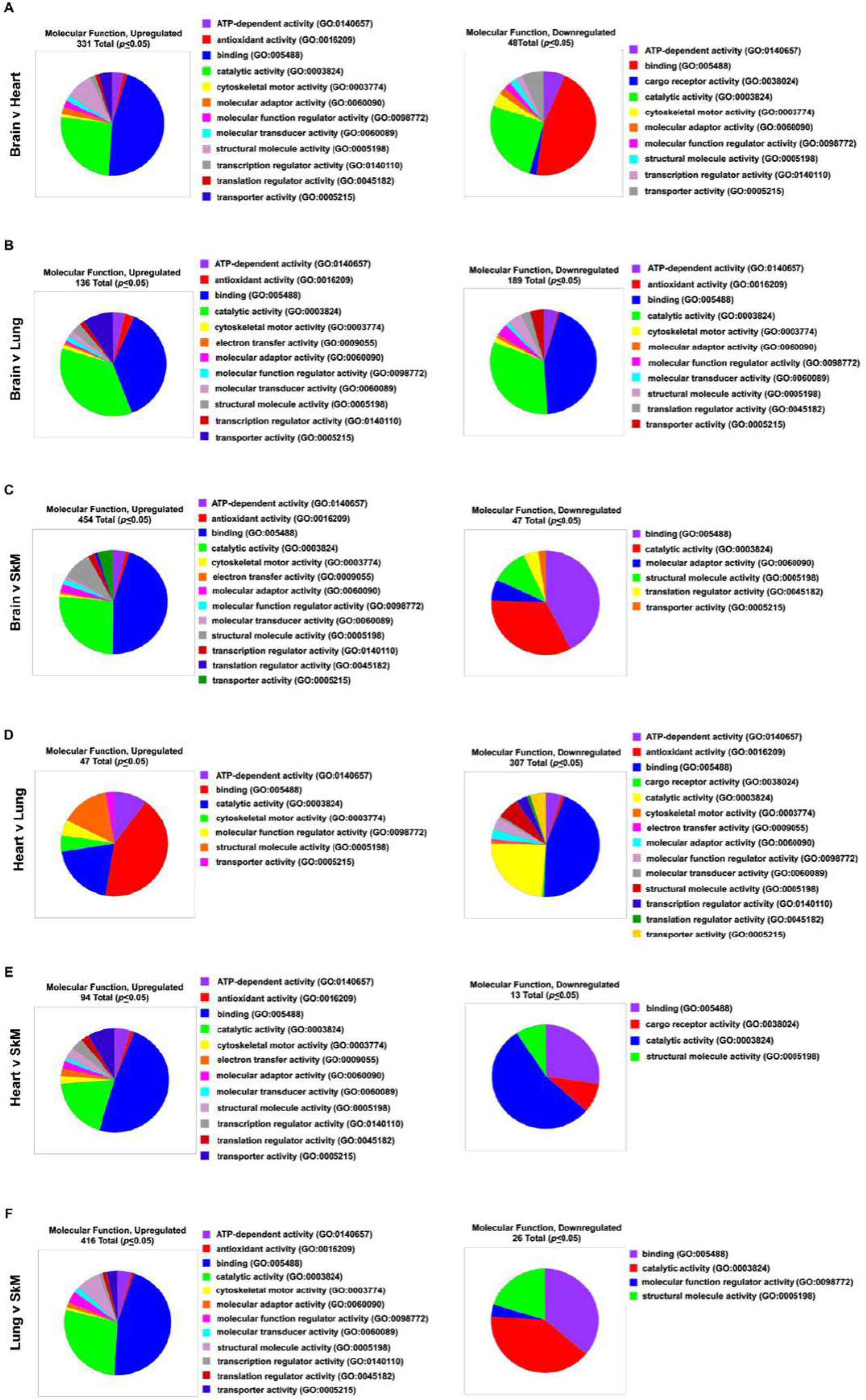
Panther Ontology for EC proteomes between tissues. Differences in the EC proteomes from Brain, Heart, Lung, and Skeletal Muscle (SkM), with target Ontology identified between EC tissues of origin.

## Notes

### Competing Interest Statement

The authors have declared no competing interest.

